# Breeding in an agricultural landscape: conservation actions increase nest survival in a ground-nesting bird

**DOI:** 10.1101/2022.10.22.513330

**Authors:** Ádám Kiss, Zsolt Végvári, Vojtěch Kubelka, Ákos Monoki, István Kapocsi, Szilvia Gőri, Tamás Székely

## Abstract

Agricultural intensification has affected wildlife across Europe, usually prompting steep declines and regional extinctions in farmland birds. Effective conservation activities are essential for preservation of biodiversity in agricultural landscape, but despite the efforts, the halting (or reversing) the decline of farmland species are still rare. Here we investigate a ground-nesting shorebird, the Collared Pratincole (*Glareola pratincola)* that has switched its habitat preferences in Central Europe in the last 20 years from alkaline grasslands to intensively managed agricultural landscapes. We show that nest success was different between three habitat types, with the highest nest success in fallow lands whereas nests in row crops showed the lowest hatching success. Nest success was also associated with timing of breeding and breeding density, since nests produced early in the breeding season and those in dense breeding sites hatched more successfully than those later in the season and low breeding density, respectively. Importantly, since 2012 direct conservation measures have been implemented that include the marking of nests and negotiating with farmers to avoid the usage of agricultural machinery around the marked areas, controlling nest predators and most recently creating suitable nesting sites and foraging areas for the Pratincoles. Due to these direct conservation actions, the probability of both nest survival increased from 0.11 in 2012 to 0.83 in year 2021, and the size of breeding population increased from 16 pairs in 2013 to 56 in 2021. Taken together, agricultural areas can continue providing important habitats for various organisms, and with targeted conservation actions we can reduce or even halt the decline of farmland species.

## Introduction

Natural habitats are disappearing or degrading at global scales at an unprecedented rate, which is a result of the combined effects of current climatic processes and land use change during the Anthropocene (Fahrig, 1997; Balmer & Erhardt, 2000; Davidson, 2014; Hu et al., 2017). One of the main reasons is the expansion of intensive forms of agricultural land-use that has led to negative changes or the complete disappearance of various habitats across Europe (O’Connor & Shrubb, 1986; Potter, 1997). These declines are especially severe in grassland (or steppe) breeding animals, considered as highly sensitive for environmental changes, as many species declined dramatically during the past decades (Fuller, 2000; Massa & La Mantia, 2010; Ward et al. 2010; Guerrero et al., 2012). As a consequence of the loss of grassland habitats, birds that traditionally bred in open natural habitats, increasingly occupy arable lands and cultivated areas during reproduction (Galbraith, 1987; Böhning-Gaese & Bauer, 1996; Brady & Flather, 1998).

However, breeding in agricultural landscapes may be costly, as these habitats may not be productive enough due to the mal-assessment of the habitat by prospective breeders (Székely 1992), and thus serve as ecological traps (Schlaepfer, et al., 2002; Robertson & Hutto, 2006; Pärt et al., 2007; Gilroy et al., 2011; Hollander et al., 2017). Additionally, the intensification of agricultural practices can affect nesting success of ground-breeding species in numerous ways, including direct elimination of nests, chicks and/or adults by mowing, cultivating by agricultural machineries, use of pesticides, irrigation, or drainage (Berg et al., 1992; Wilson et al. 2005; Kentie et al., 2013). Unfortunately, there are abundant examples showing the negative impacts of agriculture, with local species extinction from extensive areas that are full or occasional breeders of agricultural habitats including the Great Bustard (*Otis tarda*) and Grey Partridge (*Perdix perdix*) (Donald et al., 2001; Arroyo, et al., 2002; De Leo et al. 2004; Alonso & Palacín, 2010; Potts, 2012; Gooch et al. 2015). Furthermore, these pressures have been intensified owing to global climatic changes, coupled with increased predation rates in human-modified habitats. Specifically, environmental changes have boosted the populations of meso-predators which have further impacted the nest or offspring survival rates of ground-breeding birds (Roodbergen et al. 2012; Kentie et al., 2015; Kubelka et al., 2018; Brzeziński et al., 2020). To mitigate these negative effects, targeted conservation actions are needed (Arroyo et al., 2002; Zamečník et al, 2008; Schekkerman et al., 2008).

Here we report the results of a 10-years conservation effort focused on the Collared Pratincole (*Glareola pratincola*) that is affected by habitat alterations and has undergone a population decline across many parts of Europe (Yuri et al., 2020). The Collared Pratincole is a ground-nesting shorebird that historically bred in loose colonies in alkaline grasslands close to wetlands in Central Europe (Cramp and Simmons, 1983). The largest inland breeding population in the Carpathian Basin existed during the early 1900s (Aradi 1979; Kiss et al., 2018). Collared Pratincoles feed on flying insects including dragonflies, flies and various-sized *Coleoptera* species, and they build their nest into a hoofprint or on bare ground (Beretzk, 1954; Cramp and Simmons, 1983). The global population is declining (IUCN Red List, 2017) although it has been hard to assess the change of abundances of the species due to their high dispersal propensity and semi-nomadic strategies, leading to high fluctuations in breeding densities (Yuri et al., 2020). Collared Pratincoles have been shown to use agricultural lands for breeding in several parts of Europe in the past centuries (Calvo & Alberto, 1990; Calvo, 1994; Calvo & Furness, 1995; Lebedeva, 1998; Kiss et al., 2017; EBBA 2, 2020), and recently most breeding attempt occurs on arable farmland (Nardelli et al., 2015; Kiss et al., 2017; Vincent-Martin, 2007). During the past decade, the Hungarian population has fluctuated between 22 and 65 pairs, and it split between two regular breeding sites in the Nagykunság and Kiskunság regions. These breeding sites became the last remaining breeding locations for the species within the Carpathian basin (Kiss et al., 2018).

We had four objectives in our study. First, to quantify Collared Pratincole nest success and investigate the ecological and behavioural variables that may predict nest success including habitat type, timing of breeding, proximity to open water surfaces as a proxy for water availability and breeding density. Second, to compare nest survival rates between different agricultural habitats. Thirdly, to investigate the effects of conservation measures on nest survival, and finally to investigate potential associations between predator control and nest survival.

## Methods

### Study area

Data collection and conservation activities were carried out in the Nagykunság region, located in the middle of the Hungarian Great Plain, Eastern Hungary (N47.2, E20.9, Figure 1). The climate is eastern continental, characterised by dry and warm periods during the breeding season, interspersed with short, heavy rainfalls of 20 – 100 mm/hour (Hungarian Meteorological Service, 2021). We focused on the southern part of the region, where the landscape is dominated by cultivated lands, primarily rice fields (Plate 1). Due to the requirements of rice cultivation, each year more than 1500 hectares of farmland is artificially flooded, providing important habitats for breeding and migrating shorebirds (Monoki & Kiss, 2017).

**Figure 1.**
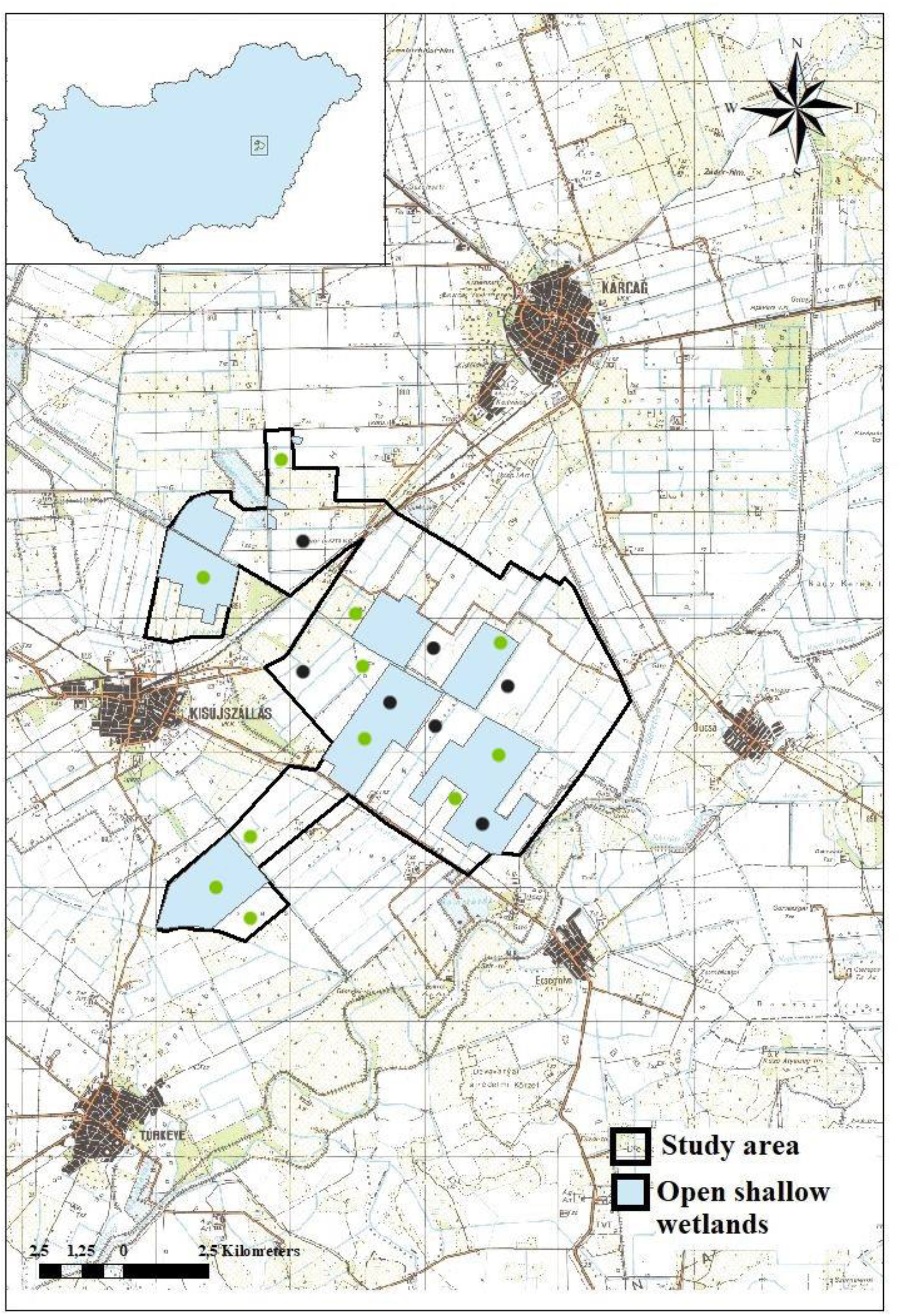
A map of the study area (12 500 hectares, black solid line), and paddy fields (blue polygons). Black dots = standard, green=alternative breeding sites.

The estimated number of breeding pairs of Collared Pratincoles fluctuated strongly between 13 and 65 pairs, between 2012 and 2021 (33.25 ± 14.61) (database of the Hortobágy National Park Directorate).

### Data collection

Starting in 2012, we continuously collected Collared Pratincole field observation records for breeding sites, including nest-site selection, nest success and behaviour. We recorded these data in field using a handheld Trimble Personal Digital Assistant, which were later processed in Arcmap 10.1. Data were also collected in i) croplands used by Collared Pratincoles and ii) shallow wetlands. Using nest points, and different agricultural variables we created a map of nest points, shapefiles of arable lands and shallow water bodies.

Nest locations, polygons of croplands and water bodies were prepared for further analyses using ArcMap 1.0 software. Monitoring activities of the potential nesting sites, localization and revisiting of nests were assisted by high-quality binoculars and spotting scopes. Nests were always approached to a distance of 8-10 meters by 4WD cars – even within agricultural fields – to avoid any disturbance of incubating shorebirds, allowing the observation of Collared Pratincole behaviour. In addition to the location of nests, we identified the type of land used for nesting, which were classified into three categories based on preparation and management technologies (Supplementary Table 1, Figure 2). After locating the nests, we recorded clutch size, nest cover, GPS coordinates for each nest, which we completed by positioning a small wooden twig 1 meter apart from each nest to be able to relocate them. After finding each new nest or colony, we consulted the owner or manager of the land. To prevent nest destruction by farming activities, we marked a buffer zone using 1.5 m tall, 2 cm thick wooden poles around the nest. However, to prevent predators from learning these signals, we only placed these markers during active agricultural work (based on a method of Zamečník et al. 2018). The mean size of these oval-shaped buffer zones amounted to 0.01 hectares, which is of sufficient size to ensure adequate protection of the nest from agricultural machinery (Figure 3).

**Figure 2.**
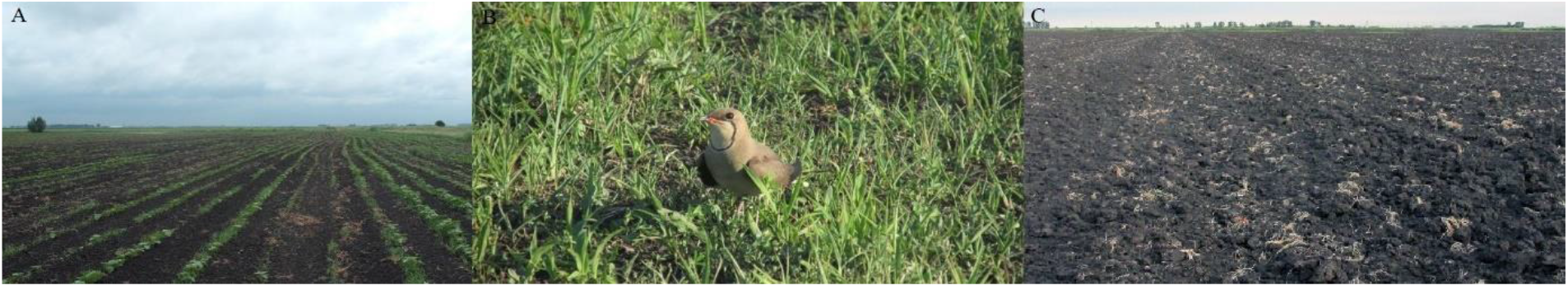
Agricultural habitats, (A) Row crop, (B) Spring-cover crop, (C) Fallow land

**Figure 3.**
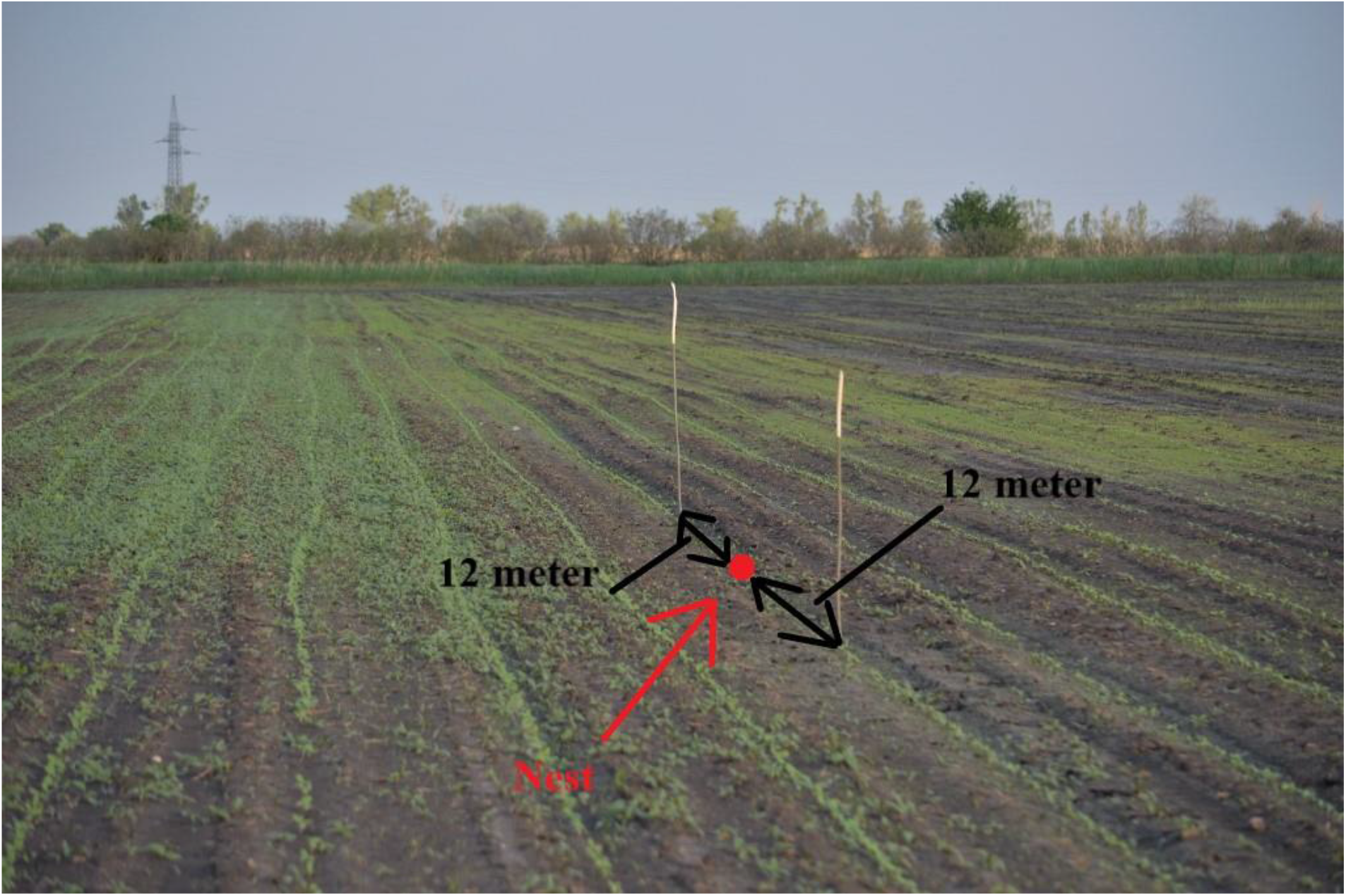
A standard-sized buffer zone indicating the nest location.

During the incubation period, all active nests were checked using spotting scope in every second day, but we revisited all nests in 2 – 3 times per week on average. After the last visit, we classified the hatching success of the nests as i) hatched; ii) predated; iii) abandoned; iv) unknown. v) flooded; vi) destroyed by agricultural machineries. To identify the fate of each nest, we used Green’s methods (1987) in addition to our field observations (Kiss et al., 2018). Trail cameras were applied for 116 nest to identify the species of nest predators and the date of hatching.

Survival and productivity of ground-nesting birds is influenced by predation (Martin, 1993; Rodbergen et al., 2012; Kubelka et al., 2018). To investigate relationships between breeding success and the number of huntable predators killed by professional hunters in the study area, we collected data from hunters. Predatory mammals include Red Fox (*Vulpes vulpes*), Golden Jackal (*Canis aureus*) and Europen Badger (*Meles meles*) and birds are represented by European Magpie (*Pica pica*) and Hooded Crow (*Corvus cornix*). We requested data from professional hunters between 2017 and 2021, and we aggregated the numbers of shot individuals for each year. Data on hunting activities were collected from the ca. 26,000 hectares large regional hunting districts which overlapped with more than 65% of the breeding sites (National Game Management Database, 2022).

### Estimating daily and total nest survival

To investigate the effects of years and agricultural habitats on nesting success, we estimated daily and total nest survival rates using Mayfield’s calculations (1975). The Mayfield’s method (1975) is applied to estimate the chances of a clutch surviving the daily and full nesting period by defining daily nest survival rate as the number of failed nests divided by the sum of exposure days. Total nest survival was calculated using Mayfield’s formula: daily nest survival ^nesting period in days^. The computation of nest survival rates requires information on total exposure time, which was defined as the number of days from finding to the day of confirmed or expected day of final fate of the nest. For all nests where signs of hatching were observed, the exposure time was calculated from the day of finding to the confirmed or predicted date of hatching. For those nests which has become depredated, this interval was calculated as starting from the day of finding until the midpoint between the last positive and the first negative visit to the nest. For all other outcomes (unknown, abandoned, flooded, destroyed by agricultural machineries), the exposure time was defined from the day of finding until the last positive visit, following the standard protocols (e.g. Kubelka et al., 2018).

### Statistical analyses

To identify relationships among individual-level reproduction success metrics, i) nests which hatched or failed and ii) number of hatched chicks, we performed Generalised Linear Models (GLM), entering year, habitat type, julian day of egg-laying start as well as distance to the a) nearest field edge; b) closest water body and c) mean distance of the three closest nests within the same colony as fixed factors. As the nest success response (hatched or failed) is a binary variable, we conducted a logistic regression-type GLM, applying ‘logit’ link error function. Field boundaries and wetlands were available in shape file formats, using our own field mappings. The mean distances of the three neighbouring nests were computed applying the ‘nndist’ spatial neighbourhood function available in the ‘spatstat’ package for spatial statistics. Egg-laying start was defined as the julian day of the record, defined as the number of days counted from 1 January each year, considering the first day for incubation for individual clutches.

Further, we analysed the relationship between i) clutch size and habitat type; ii) timing of hatching and habitat type iii) daily nest survival metric (aggregated for years and habitats) based on the number of culled predators by official hunters, applying ANOVA-tests implemented using the ‘lm’ function. All statistical analyses were performed within the R v. 3.3.3 statistical programming environment (R Core Team, 2021).

## Results

### Breeding success and timing of breeding

The breeding season of Collared Pratincoles lasted from late April to early August. The first eggs have hatched on 16 May, which implies that the clutch was completed on 29 April. The latest hatch date was recorded on 3th August. The mean egg-hatching date was found on 15 June ± 1.2 days over the study period, and most nests were produced between 25 May and 15 June (n = 212 nests, Supplementary Figure 2).

Collared Pratincoles bred in three types of habitats. The majority of nests were found in row crops (48%), followed by fallow lands (29%) and spring cover crops (23%) (total n = 315 nests, Table 2). Timing of breeding was different between crop types: nests in row crops or spring-cover crops hatched earlier than in fallow lands (One-way ANOVA, b > 6.311, F_2,210_ = 39.02, p_min_ = 0.009, Supplementary Figure 1). Similarly, clutch size was significantly related to habitat type: the largest clutch sizes were found in row crops (One-way ANOVA, b < -0.0491, F_2_,_268_ = 2.601, p_min_ = 0.0254). The number of successful hatchings per all nesting in different agricultural habitats was 74.7% (n = 68) in fallow lands, 69.4% (n = 50) on spring cover crops and 61.8% (n = 94) in row crops. The total number of clutches hatched successfully was 67.3% (n = 212 nests).

**Table 1.**
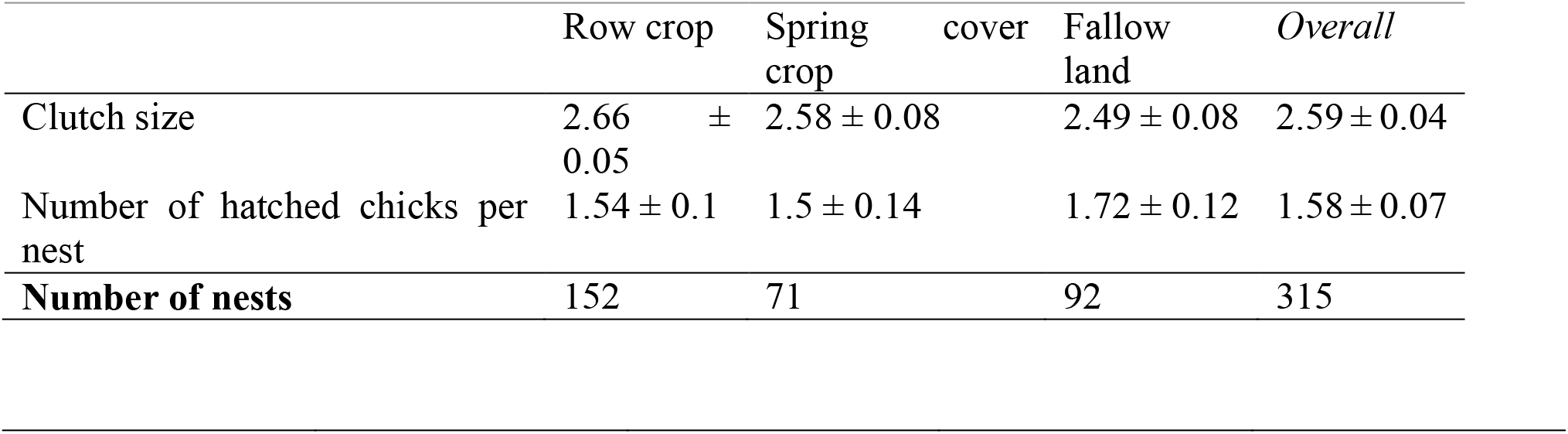
Clutch size and nest-success in agricultural habitats of Collared Pratincoles in Hungary (mean ± SE).

**Table 2.** Relationships between hatching success (A.) and the number of hatched chicks (B.), fitted by a logistic and linear regression analyses (GLM) as a function of agro-technology, time, and space, and ecological variables. Significant relationships are indicated in bold.

| Variables | A. |  |  |  | B. |  |  |  |
| --- | --- | --- | --- | --- | --- | --- | --- | --- |
|  | Esti-<br>mate | SE | z-<br>value | p-value | Esti-<br>mate | SE | t-<br>value | p-value |
| Intercept | -463.119 | 124.423 | -3.722 | < **0.001** | -292.035 | 63.835 | -4.575 | < **0.001** |
| **Agricultural** |  |  |  |  |  |  |  |  |
| Spring-cover crops | 1.032 | 0.403 | 2.560 | **0.010** | 0.423 | 0.200 | 2.114 | **0.035** |
| Fallow lands | 1.354 | 0.447 | 3.029 | **0.002** | 0.719 | 0.216 | 3.321 | **0.001** |
| **Ecology and timing** |  |  |  |  |  |  |  |  |
| Year | 0.231 | 0.062 | 3.741 | < **0.001** | 0.146 | 0.032 | 4.624 | < **0.001** |
| Egg-laying start | -0.012 | 0.009 | -1.357 | 0.175 | -0.011 | 0.005 | -2.285 | **0.023** |
| Field boundary | 0.003 | 0.003 | 0.954 | 0.340 | 0.001 | 0.001 | 1.006 | 0.316 |
| Distance to water<br>body | < 0.001 | < 0.001 | 0.403 | 0.687 | < 0.001 | < 0.001 | 1.559 | 0.120 |
| **Social behaviour** |  |  |  |  |  |  |  |  |
| Breeding density | < 0.001 | < 0.001 | -1.391 | 0.164 | < 0.001 | < 0.001 | -2.468 | **0.014** |

Nest success was related to the type of habitat: pairs that chose spring-cover crops or fallows showed higher nest success and more hatched chicks, compared to pairs that bred on croplands. In addition to crop type, nest-success was associated with time in the season and breeding density, since early nests and those that had higher breeding densities produced more chicks. (Table 2).

### Nest survival and causes of nest-failure

Nest failures were caused by predation (56.7%), nest abandonment (23.1%), flooding by heavy rainfalls (18.3%), and unknown (0.9%) (total n = 104 nests). As a result of the nest-marking scheme, agricultural machinery destroyed relatively few nests (0.9%) (Supplementary table 2). 83.1% of all nest predation (n = 49) and 89.5% (n = 17) of all flooded nests were found in row crops and spring cover crops. The most common nest and fledgling predators included mammalian predators that include Red Fox, European Badger and birds, especially Hooded Crow, Western Marsh Harrier (*Circus aeruginosus*), and Caspian Gull (*Larus cachinnans*).

Daily nest survival significantly increased over the study period (linear regression, b= 0.0064, N=8, p = 0.0189, Figure 4), but there was no significant association found between habitat type and daily survival (Two-way ANOVA, b= -0.0457, SE = ± 0.0355, p-value = 0.209) or total nest survival (Two-way ANOVA, b= -0.0917, SE = ± 0.1592, p-value = 0.569). The highest level of total nest survival was recorded for pairs nesting in fallow lands (Supplementary table 3).

**Figure 4.**
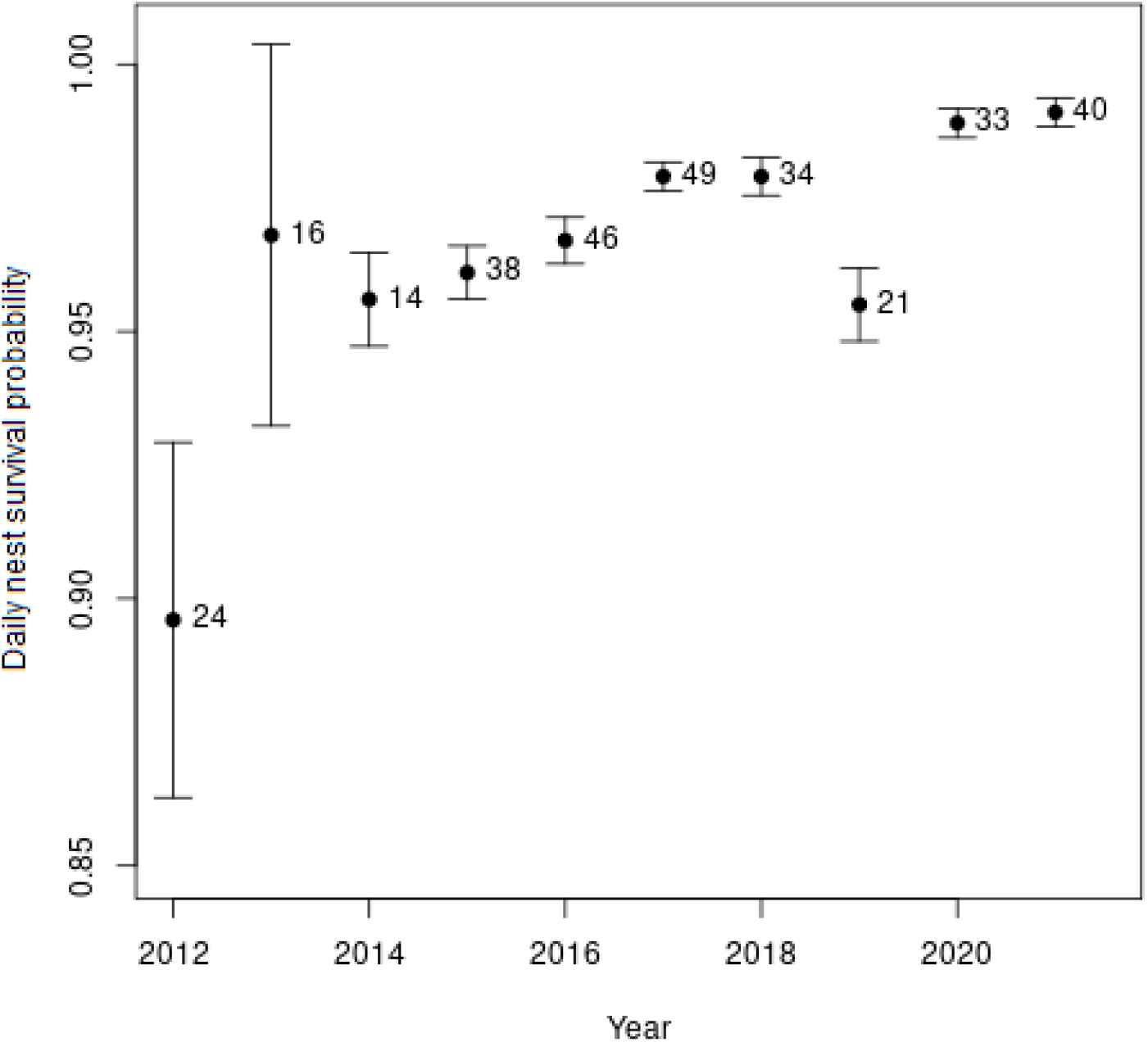
Daily nest survival (mean ± SE) and the number of nests (in parentheses) during the study years (r^2^=0.5482, n = XX years).

### Conservation action

Daily nest survival was not predicted by either the number of avian predators (linear regression, b = 0.0001, N = 5, p = 0.655) not by the number of mammalian predators (linear regression, b = 0.0005, N = 5, p = 0.503); however, we had data only for five years. The hunting pressure increased over the study period as indicated by the increasing number of removed predators between 2017 and 2021. (Supplementary material Table 4).

During the study period, we needed to carry out direct conservation interventions in the form of a buffer-zone designation at 159 nests (50%, n = 315). The number of directly protected nests fluctuated among the years and habitats, although the largest proportions of nests which had to be protected were located in croplands (92% n = 159, Table 3).

**Table 3.**
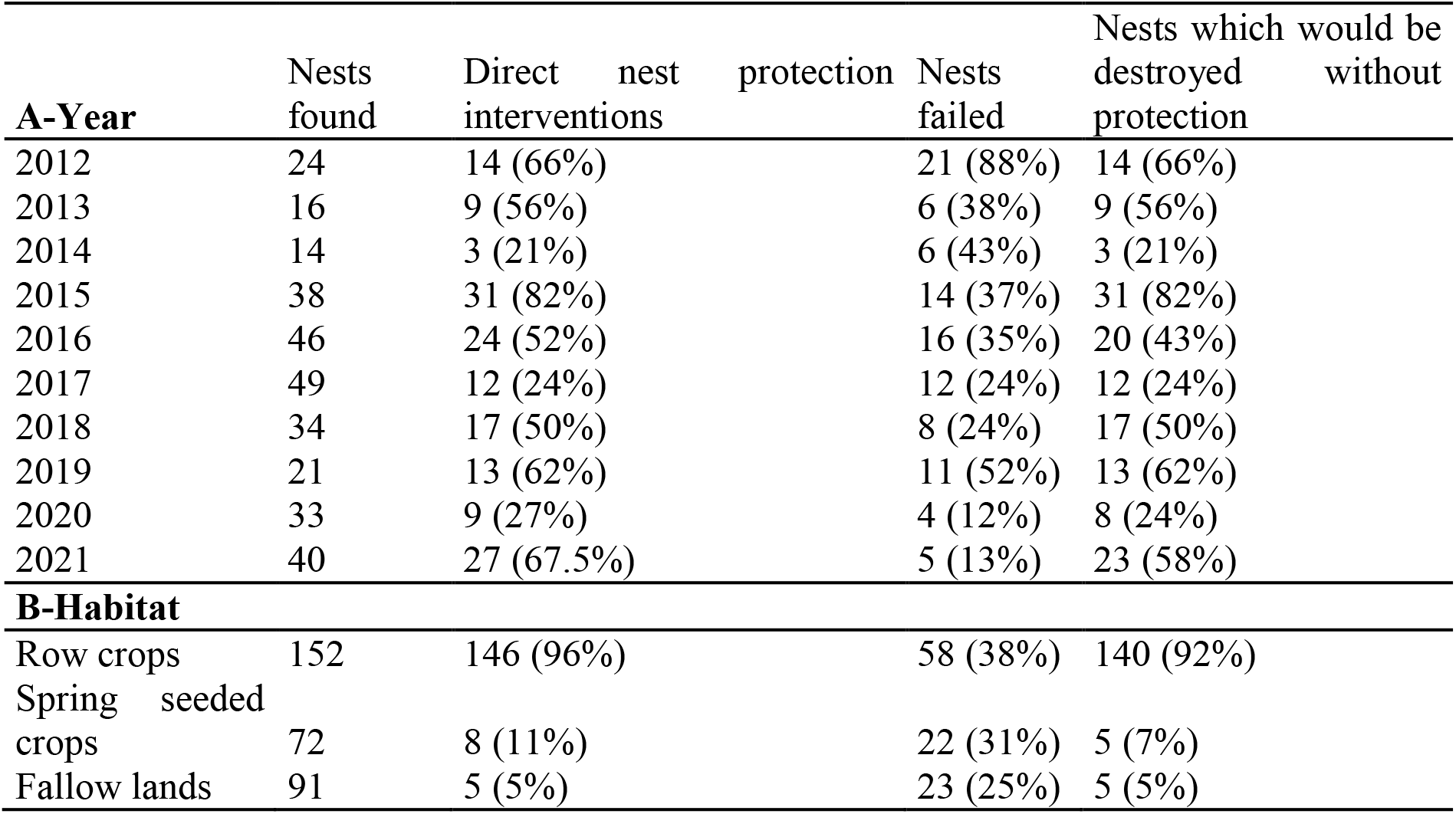
Relationships between direct conservation efforts and agricultural habitat types (n=315 nests).

## Discussion

Our key findings are that i) habitat and the density of colony correlated with nest-success, ii) nest success, and daily survival rate increased constantly over the years and iii) survival rates showed similar levels for nests found in row crops as the other two habitat types, where large-scale protection zones had to be established to protect them.

Maintaining good relationships between farmers and conservationists is essential to achieve success of conservation projects (Logsdon et al., 2015; Homberger et al., 2017). Similarly to our study species, many other shorebirds have experienced a decline in their optimal breeding habitats across Europe, and thus they have been forced to choose riskier breeding sites in terms of survival chances as majority of traditional habitats were converted for agricultural use (Berg, 1992; Schifferli et al., 2006; Kentie et al., 2015). In the case of Collared Pratincole, some studies compared the artificial and natural habitats in terms of nest-site selection and survival (El Malki et al., 2013) and found higher nesting success in natural habitats (Calvo, 1994; Vincent-Martin, 2007). In our study area, the species only occupied agricultural habitats, which allowed us to compare various habitats that were created using several types of agricultural treatments as well as various types of soil structures and vegetation cover. Most fields were used in different phases during the breeding season of the species, therefore the peak of hatching dates were observed at different times. Nesting strategies were highly dependent on the local agricultural schedule because row crops and spring cover crops were sown first and were followed by the ploughing of fallow grounds. In row crops and spring-sown fields, vegetation grows particularly uniformly and rapidly, reducing the time period in which Collared Pratincoles are able to nest successfully. By contrast, vegetation on fallow land grows heterogeneously in mosaic spots, creating better nesting conditions. The highest number of nests was found in row crops, which was the most abundant type of agricultural breeding habitat available for Collared Pratincoles and other shorebirds in any given year. Average clutch size was similar to those found in Mediterranean areas such as in Spain (Bertolero & Martinez-Vilalta, 1999) and France (Vincent-Martin, 2007), and higher than those breeding in northern coastal areas of the Azovian Sea (Pozhidaeva & Molodan, 1992) and in Moroccan coastal wetlands (Elmalki et al., 2013). Average number of hatched chicks were lower than in the coastal habitats of the Azov Sea (Pozhidaeva & Molodan, 1992) and in Algeria (Bensaci et al., 2014).

We found that Collared Pratincoles that chose fallow lands and spring-cover crops had significantly higher nest-success during breeding season than those nesting in row crops. Nest survival rates were also influenced strongly by the timing and intensity of agricultural operations for other shorebirds. For example, in the case of Northern Lapwing *(Vanellus vanellus)*, nest losses depended on the timing of spring tillage during the nesting period but was independent of crop type (Sheldon et al., 2007). We observed that predation pressure was lower in extensively used habitats, as compared with intensively treated areas. Similar patterns have been documented in Black-tailed Godwit *(Limosa limosa)* (Kentie et al., 2015) and other ground-nesting species. It is likely that the rise of modern intensive agriculture has favoured generalist predators by providing opportunities to colonise more readily in the riskier breeding sites (Pescador & Peris, 2011). In addition, large amounts of rapid rainfall (> 20 mm / hour) were less likely to flood nests in fallow fields, as the repeated compaction of soil during agricultural management reduces the ability of water to freely drain.

Collared Pratincole frequently breed in colonies with various sizes, thus inter-nest distance was expected to be an important predictor of nest success. In the case of a similar species, the Pied Avocet (*Recurvirostra avosetta)*, nesting success was lower in both denser and less dense colonies, and higher at intermediate densities, as observed by Hötker (2000). The number of breeding Collared Pratincoles in Hungary is significantly smaller in comparison to other populations in Europe, so these effects presumably did not apply.

We found no difference between either daily and total survival rate of nests and habitat type. Nests threatened by agricultural machinery (mainly in row crops) were effectively protected, and thus intensive conservation activities buffered the difference in nest survival rates between different habitats. Nest survival rates could have been influenced by predation and heavy rainfalls between habitats, but these effects were quite low over the years. Direct nest protection interventions allowed the spectacular increase of the level of survival rates. Similar effects were experienced in the Wood Turtle (*Glyptemys insculpta)* conservation project, which found that nest success can be increased spectacularly applying adequately designed interventions (Bougie et al., 2020). In the absence of direct nest protection, breeding success would have also been low in critical habitats, similar to that described by Calvo (1994), who found that as a result of changing agricultural practices, nesting success could also improve. Nest-success and daily survival rate of nests has increased significantly over the past decade, probably due to the qualitative and practical development of nest search, and conservation practice. Moreover, the intensity of agricultural practice has noticeably decreased in Hungary in the past decades (Báldi & Batáry, 2011).

### Conservation activities

Since the Collared Pratincoles abandoned their traditional breeding habitats in alkaline grasslands, the Hortobágy National Park Directorate has made several attempts to improve their natural habitats in order to re-establish the species’ breeding populations, so far without success, but see (Kovács & Kapocsi, 2005). In addition to restoration efforts, the national park also carried out a parallel search and protection of colonies and solitary pairs nesting in their active breeding sites. Accordingly, these breeding grounds are managed by an intensive agricultural land use scheme, thus the system of conservation management requires a composed and precise cooperation between conservationists and farmers (Kiss et al., 2018). Direct nest protection activities had to be implemented mostly in row crops, because these types of agricultural land (especially sunflowers, corn fields) are cultivated intensively during the breeding season. In contrast, on spring-cover crops and fallow lands – with a few exceptions – we detected no disturbance by agricultural machineries after ploughing or seeding. Although we didn’t find a significant correlation between the numbers of shot predators and the daily or total survival rates of nests, this probably as a result of inaccurate data and small sample size, we chose to continue managing this activity, as targeted lethal and non-lethal predator removal programs are important for long-term conservation of ground-nesting bird species, especially shorebirds, proven by successful conservation programmes for many species (Neuman et al., 2004; Bolton et al., 2007). It is likely that there was a positive change in the efforts of predator hunting (increase in the number of hunting days, and growing level of efforts to hunt game predators), although no written resources, but only verbal information was available to explain this pattern. Thus, a more thorough investigation into the relationships among predator removal and breeding success of Collared Pratincole in the coming years.

We did not experience significant nest mortality by predators in nests marked with poles as opposed to unmarked nests, similar to that observed by Zámeĉník et al. (2018), as poles were left near nests only during critical, endangered periods. Thanks to the positive attitude of local farmers, the organization of protection was feasible and effective, and none of the known nests were at serious risk during agricultural work. However, the long-term protection of Collared Pratincoles should be further improved by the establishment of fields and fallows free from agricultural disturbance. As a result of the current agricultural scheme, agricultural land in Hungary and Europe is typically used too intensively, arable fields typically do not remain without crops for a significant part of the year (Tarjuelo et al., 2020).

On global scales, human activity can negatively influence the behaviour, productivity, and nest survival of ground-nesting birds in various habitats, especially on farmlands (Fahrig, 1997; Donald et al., 2001; Colwell, 2010, Ward et al., 2010). We have set ourselves the goal of habitat development at the local level, as a result of which 50-100 hectares of fallow land are created every year to facilitate the settlement of shorebirds on the Nagykunság rice systems. These barren fields are created by disc-ploughing during the end of April, and after the treatment there is no human disturbance during the breeding season. These areas are considered insignificant in relation to the size of the total habitat, but this seems to be a promising project as a variable number of birds have nested and gathered in these fallow areas in recent years. An improved solution could be supported by the development of targeted agricultural programs, which would specifically set management standards for the arable land used by the species, and also provide financial support to farmers’ efforts.

Taken together, our results suggest that it is worth maintaining intensive conservation activities to protect rare or endangered species, as we can achieve success even in intensively managed habitats. Without such interventions, large proportions of farmland bird nests are lost to agricultural machinery and the remaining Eurasian population of Collared Pratincole out of Hungary might be considerably threatened by anthropogenic pressure. In addition to effective direct nest protection, it is important to increase the proportion of safe fallow lands in the future as a specific agri-environmental protection measure so that as many farmland birds as possible have the opportunity to choose this undisturbed agricultural habitat for breeding. As Collared Pratincoles nest in several places in artificial habitats in Europe, mainly close to secondary wetlands, such as rice fields, the breeding habitat protection or restoration should be more widely implemented to maintain biodiversity in agricultural landscapes.

## Author contributions

Study design and fieldwork: ÁK, ÁM, IK, TSz, SzG; data analysis and writing the article; ÁK, ZsV, VK, ÁM, IK, SzG, TSz.

## Conflict of interest

None

## Ethical standards

None

## Acknowledgements

We thank Antal Széll, Miklós Lóránt and other national park rangers who participated in the conservation management and data collection of the species in Hungary. We are grateful to Fanni Takács who supported our field data collection, and we are also grateful to William Jones, who provided linguistic assistance and commented on the earlier draft of the manuscript. We thank the farmers of Kisújszállás and Karcag especially to the workers of Nagykun 2000, Hubai és Társa, Indián Rizs Agro Inc. who supported the protection of the species in Hungary with their patience and attitude and helping at the agricultural executions. We also thank the professional hunters for providing the planned predator control. V.K. and T.S. were supported by ÉLVONAL-KKP 126949 of the Hungarian government, and Eotvos Lorand Research Network (Grant no. 1102207).

## Notes

### Competing Interest Statement

The authors have declared no competing interest.

