## Supplementary Table 1 for "Breeding in an agricultural landscape: conservation actions increase nest survival in a ground-nesting bird"

### Supplementary Table 1. Agricultural nesting site categories of Collared Pratincole *Glareola pratincola* in Hungary.

| Habitat | Characteristics of preparation and treatment | Examples |
| --- | --- | --- |
| Row crops | Planted in wide rows, intesive land use with pesticides and row cultivation | corn, sorghum, sunflower |
| Spring cover crops | Planted in narrow rows, less intense land use by chemicalization | Alexandra-clover, alfalfa, barley, millet, oat, phacelia |
| Fallow lands | Created by disc ploughing, there aren’t any disturbance by agricultural machineries | dry paddies, rested areas |

Supplementary Table 2. Nest fates, exposure days, daily and total nest survival rate probabilites of Collared Pratincoles.

| Year | Hatched (%) | Predated (%) | Abandoned (%) | Flooded (%) | Unknown (%) | Agricultural activities (%) | Exposure (days) | Daily nest survival rate | Total nest survival rate (%) |
| --- | --- | --- | --- | --- | --- | --- | --- | --- | --- |
| 2012 | 13 | 71 | 0 | 17 | 0 | 0 | 202.5 | 0.896 | 0.6 |
| 2013 | 68 | 12 | 14 | 6 | 0 | 0 | 189 | 0.974 | 27.9 |
| 2014 | 57 | 14 | 7 | 22 | 0 | 0 | 136 | 0.956 | 11.7 |
| 2015 | 63 | 26 | 11 | 0 | 0 | 0 | 363.5 | 0.961 | 15.5 |
| 2016 | 66 | 9 | 4 | 17 | 2 | 2 | 478 | 0.967 | 19.8 |
| 2017 | 76 | 16 | 8 | 0 | 0 | 0 | 616 | 0.981 | 39.3 |
| 2018 | 76 | 21 | 3 | 0 | 0 | 0 | 387 | 0.979 | 37.1 |
| 2019 | 48 | 24 | 19 | 9 | 0 | 0 | 246 | 0.955 | 11.4 |
| 2020 | 88 | 3 | 9 | 0 | 0 | 0 | 371.5 | 0.989 | 59.8 |
| 2021 | 88 | 7 | 5 | 0 | 0 | 0 | 557 | 0.991 | 65.2 |

Supplementary Table 3. Number of hatched nests, proportion of exposure days, daily and total nest survival probabilities during the study period.

|  | Row crops | | | | | | Spring cover crops | | | | Fallow lands | | | | | |
| --- | --- | --- | --- | --- | --- | --- | --- | --- | --- | --- | --- | --- | --- | --- | --- | --- |
|  | **No. Nests** | **No. of hatched nests** | **Exposure (days)** | **Daily nest survival** | **Total nest survival rate (%)** | **No. nests** | **No. of hatched nests** | **Exposure (days)** | **Daily nest survival** | **Total nest survival rate (%)** | | **No. nests** | **No. of hatched nests** | **Exposure (days)** | **Daily nest survival** | **Total nest survival rate (%)** |
| 2012 | 14 | 0 | 139.5 | 0.890 | 0.390 | 7 | 2 | 33 | 0.848 | < 0.001 | | 3 | 1 | 30 | 0.930 | 3.1 |
| 2013 | 9 | 5 | 142.5 | 0.972 | 25.90 | 3 | 2 | 40.5 | 0.950 | 8.7 | | 4 | 4 | 6 | 1 | 100 |
| 2014 | 3 | 0 | 10 | 0.7 | < 0.001 | 6 | 3 | 47 | 0.936 | 4.3 | | 5 | 5 | 79 | 1 | 100 |
| 2015 | 21 | 10 | 216.5 | 0.949 | 8.3 | 8 | 7 | 90.5 | 0.989 | 59.1 | | 9 | 7 | 56.5 | 0.965 | 18.4 |
| 2016 | 24 | 18 | 309 | 0.981 | 40.2 | 15 | 7 | 99 | 0.920 | 1.9 | | 7 | 5 | 70 | 0.971 | 24.7 |
| 2017 | 12 | 8 | 170.5 | 0.976 | 31.5 | 10 | 9 | 170.5 | 0.994 | 75.1 | | 27 | 20 | 274 | 0.974 | 28.6 |
| 2018 | 17 | 12 | 215.5 | 0.977 | 33.1 | 2 | 2 | 36 | 1 | 100 | | 15 | 12 | 135.5 | 0.980 | 38.3 |
| 2019 | 17 | 8 | 192.5 | 0.950 | 8.7 | 1 | 0 | 3 | 0.670 | < 0.001 | | 3 | 2 | 50.5 | 0.981 | 40.2 |
| 2020 | 9 | 9 | 136 | 1 | 100 | 16 | 15 | 180.5 | 0.995 | 78.8 | | 8 | 5 | 50 | 0.940 | 5.3 |
| 2021 | 26 | 24 | 356 | 0.994 | 75.1 | 4 | 3 | 32.5 | 0.970 | 23.5 | | 10 | 8 | 168.5 | 0.988 | 56.4 |
| *MEAN DNSR+TNSR* |  |  |  | *0.940* | *32.3* |  |  |  | *0.930* | *35.2* | |  |  |  | *0.970* | *415.* |

Supplementary Table 4. Number of removed predators (individual) by official hunters from the study site between 2017 and 2022. Data are not available for years before 2017.

| Year | Mammals | Birds |
| --- | --- | --- |
| 2017 | 280 | 263 |
| 2018 | 488 | 312 |
| 2019 | 415 | 699 |
| 2020 | 504 | 1046 |
| 2021 | 582 | 953 |
| Total number of individuals | 4538 | 6546 |

Supplementary figure 1. Phenology of nest hatching events during a breeding season in various agricultural habitats (n=212).

Number of nests

| Hatching date (Five-day periods) | Row crops | Spring cover crops | Fallow lands |
| --- | --- | --- | --- |
| [6-10] May | 0 | 0 | 0 |
| [11-15] May | 0 | 0 | 0 |
| [16-20] May | 2 | 0 | 0 |
| [21-25] May | 2 | 1 | 2 |
| [26-31] May | 19 | 14 | 0 |
| [01-05] June | 24 | 11 | 2 |
| [06-10] June | 14 | 3 | 9 |
| [11-15] June | 21 | 1 | 11 |
| [16-20] June | 7 | 4 | 4 |
| [21-25] June | 4 | 1 | 5 |
| [26-30] June | 1 | 6 | 8 |
| [1-5] July | 1 | 4 | 8 |
| [6-10] July | 0 | 1 | 3 |
| [11-15] July | 0 | 0 | 10 |
| [16-20] July | 0 | 3 | 0 |
| [21-25] July | 0 | 1 | 3 |
| [25-31] July | 0 | 0 | 1 |
| [1-5] August | 0 | 0 | 1 |
